# Differential expression of mitochondria-associated genes in clinical samples of *Plasmodium falciparum* showing severe manifestations

**DOI:** 10.1101/2025.05.24.655897

**Authors:** Sukriti Gujarati, Bharat Raj Singal, Dinesh Gupta, Sanjay Kumar Kochar, Dhanpat Kumar Kochar, Ashis Das

## Abstract

The malaria parasite mitochondrial proteins are critical targets for antimalarial drugs, however, the emergence of drug resistance against the existing protein targets necessitates novel therapeutic approaches. In this study,we profiled mitochondrial sense and natural antisense transcripts (NATs) in 22 *Plasmodium falciparum* clinical isolates, classified into three disease groups - uncomplicated malaria (UNC, n=6), cerebral malaria (CM, n=4), and hepatic dysfunction (HD, n=12), using a custom, strand-aware 60K microarray, and validated the transcriptome using pooled, strand-specific RNA-seq (uncomplicated pool and complicated pool). Differential gene expression in the CM and HD cohort was obtained by comparing with uncomplicated malaria (UNC) as the control cohort. The analysis revealed distinct disease-specific sense and NATs, encoded by the mitochondrial genome, along with those encoded in the nucleus and targeted to the parasite mitochondria. Although mitochondrial activity is known to be reduced in blood-stage malaria, upregulation of genes linked to tricarboxylic-acid-cycle and electron-transport in the CM cluster indicates disrupted mitochondrial bioenergetics in severe disease. Profiling of sense and antisense mitochondrial transcripts reveal a correlated expression of sense-antisense transcript pairs in both the disease manifestations, indicating a potential regulatory role of NATs in mitochondrial function. These data provide direct evidence of NATs originating from the parasite mitochondrial genome and nominate key NATs against core mitochondrial functions as potential non-conventional antimalarial targets.

## Introduction

*Malaria* is an acute febrile illness caused by the *Plasmodium* parasite and transmitted through the bite of the female *Anopheles* mosquito. *Plasmodium* infection can be broadly classified as uncomplicated or severe. Severe malaria (SM) occurs when infections are complicated by vital organ dysfunctions or aberrant metabolism [1–2]. Human infection begins with a hepatic stage after which the parasite inhabits the red blood cells and traverse through different asexual blood stages to complete a ∼48-hour intraerythrocytic cycle that involves extensive DNA replication, endomembrane expansion and organelle biogenesis [3].

The ultrastructure of the parasite cell has revealed the division of the *Plasmodium* mitochondria from a narrow, tubular structure in early ring stages to getting extensively branched on the onset of schizogony [3–4]. Known to play a ‘minimal’ role during blood stages, *Plasmodium* mitochondrion spans a highly conserved genome of 6 kb encoding for only three protein-coding genes. The mitochondrial genome also encodes multiple discrete fragments of large and small rRNA subunits, which combine with nuclear-encoded ribosomal proteins to form mitoribosomes [5–7]. Thus, the parasite mitochondria rely significantly on the influx of proteins and RNA molecules from the nucleus for diverse biological functions, including heme and pyrimidine biosynthesis, maintenance of transmembrane potential for electron transport, and oxidative phosphorylation. Mitochondria also can replicate their DNA and transcribe it with the help of nuclear-encoded molecules. [8–9]. This diverse functionality of the parasite mitochondrion makes this organelle an attractive target for various antimalarial drugs [10–12]. However, experimental evidence of increased drug resistance against these conventional “druggable” mitochondrial protein targets has been widely reported [13–14]. This calls for new, effective antimalarial approaches that can influence disease pathogenesis by manipulating gene expression.

Advances in transcriptomics have identified non-coding RNAs, especially NATs, , for their role in manipulating gene expression, suggesting their use as potential ‘non-conventional’ drug targets [15–16]. NATs (NATs) are transcribed from the DNA strand complementary to a region harbouring a sense transcript of either protein-coding or non-coding genes, influencing the upregulation or downregulation of the sense transcripts [17–18]. The emergence of NATs and their potential role in regulating gene expression have been widely reported in the malaria parasites [19–21]. We have previously reported the presence of NATs in the nuclear transcriptome of clinical isolates of *P. falciparum* [22]. This study focuses on both sense and NATs that are either encoded or targeted to the parasite mitochondria across two severe malaria manifestations -hepatic dysfunction and cerebral malaria [23–24] and reports the *P. falciparum* mitochondria transcriptional profile.

## Materials and Methods

### Consolidating genes with potential mitochondrial localization

A systematic attempt was made to consolidate a list of *P. falciparum* mitochondrial genes using web-based prediction tools, functional annotation, and experimental validation as published in the literature. Due to the lack of a single, well-characterized mitochondrial localization signal in *Plasmodium* and the lack of a single error-free mitochondrial localization prediction tool, the nuclear-encoded and mitochondria-targeted genes were predicted using multiple web tools based on different algorithms and trained on different datasets as suggested by Sun and Habermann [25]. Four major web-based prediction tools used were -TargetP, based on the mitochondrial transit peptides (mTp) enriched with arginine, leucine, and serine, responsible for targeting proteins in the mitochondrial matrix [26], Cello2GO utilizing the BLAST homology searching approaches [27], MitoFate, that uses mitochondrial pre-sequence motifs and other classical factors for predicting mitochondrial proteome [28] and PFMpred, which primarily predicts mitochondrial proteins based on amino acid and dipeptide compositions [29]. Apart from these prediction tools, genes from PlasmoDB showing a mitochondrial targeting domain are also taken into account. The final gene list was prepared by considering their predicted occurrence from at least a consensus of two or more tools to increase the reliability of the potential mitochondria-targeted genes [30]. A comprehensive literature survey was also done to include genes experimentally validated and reported as part of the mitochondrial proteome [31–35].

### Blood sample processing

Venous blood samples were collected from *P. falciparum*-infected adult patients admitted to S.P. Medical College, Bikaner, India, and passed through a Histopaque density gradient to enrich for infected erythrocytes. Informed consent was obtained from all individual participants included in the study. The team of clinicians at S.P. Medical College, Bikaner, collected patient samples according to hospital guidelines as per approval. Infection with *P. falciparum* was confirmed by slide microscopy and RDTs. Mono-infection by *P. falciparum* was further confirmed by PCR diagnosis [36–37]. Patient samples were grouped into 3 categories, as per the WHO-defined clinical guidelines [2, 38] as tabulated in Table S1 (Supplementary Sheet 1), and hybridized individually on single-color custom-designed 8*60K microarrays.

### Microarray Hybridization

The RNA from blood samples was extracted as per standard protocols [22] and used after the concentration and purity of the RNA were evaluated using the Nanodrop Spectrophotometer (Thermo Scientific; 2000) and Qubit (Thermo Scientific, USA) assay. The integrity of the RNA was analyzed on Tapestation (Agilent). The microarray hybridization and scanning were performed at the Agilent-certified microarray facility of Genotypic Technology Pvt. Ltd., Bengaluru, India. The samples were labeled using an Agilent Quick-Amp labeling Kit (p/n5190-0442). Total RNA was reverse transcribed at 40°C using an oligo-dT primer tagged to a T7 polymerase promoter and converted to double-stranded cDNA. Synthesized double-stranded cDNA was used as the template for cRNA generation. cRNA was generated by in vitro transcription, and the dye Cy3 CTP(Agilent) was incorporated during this step. The cDNA synthesis and in vitro transcription steps were carried out at 40°C. Labeled cRNA was cleaned using Qiagen RNeasy columns (Qiagen, Cat No: 74106), and quality was assessed for yields and specific activity using the Nanodrop ND-1000. Labeled cRNA samples were fragmented at 60°C and hybridized onto an Agilent Custom *Plasmodium falciparum* Gene Expression Microarray 8X60K. Fragmentation of labeled cRNA and hybridization were done using the Gene Expression Hybridization kit (Agilent Technologies, In situ Hybridization kit, Part Number 5190-0404). Hybridization was carried out in Agilent’s SureHyb Chambers at 65°C for 16 hours. The hybridized slides were washed using Agilent Gene Expression wash buffers (Agilent Technologies, Part Number 5188-5327) and scanned using the Agilent Microarray Scanner (Agilent Technologies, Part Number G2600D). Raw data extraction from images was obtained using Agilent Feature Extraction software.

### Microarray design

60K sense-antisense microarray for *P. falciparum* was designed using annotations of the *P. falciparum* 3D7 genome from PlasmoDBv57 (www.plasmodb.org). These arrays comprise 60-mer oligonucleotide probes, covering ORFs from the nuclear, apicoplast, and mitochondrial genomes. Most ORFs had five probes each for sense and antisense orientation. These probes were checked against the human transcriptome, and all cross-hybridizing probes were removed. The probes for the plasmodial mitochondrial sequence were also scanned against the *Homo sapiens* mitochondrial genome to check for cross-hybridization. The arrays were designed and printed on the Agilent platform. Probes from our earlier validated 244K microarray have been incorporated into the current version of the 60K array [22]. Here, we report only the transcripts detected by the hybridization, which are either encoded by the mitochondrial genome itself or targeted to the mitochondria.

### Microarray Data Analysis

RNA from leukocyte-depleted blood samples from 22 patient isolates, clinically categorized as uncomplicated malaria (UNC, n=6), hepatic dysfunction (HD, n=12), and cerebral malaria (n=4), were individually hybridized on custom-designed 8*60K Agilent microarrays. Feature-extracted raw image data, two-fold above background intensity, was obtained and analyzed using Agilent GeneSpring GX Software. The data was normalized in GeneSpring GX using the 75th percentile shift method. To make the analysis highly stringent, only the probes detecting transcripts in at least 40% of samples in each condition (including complicated and uncomplicated) were taken further into analysis. Further, probes showing uncorrelated expression for the same transcript were filtered out by calculating the Pearson Correlation Coefficient with a cutoff of >=0.8. Average fold expression values were calculated for each gene for the two test groups and differential gene expression was calculated for both the severe disease manifestations (CM vs UNC and HD vs UNC). Significant genes upregulated with fold change >=0.8 (logbase2) and downregulated with fold change <=-0.8 (logbase2) in the test samples with respect to the control group (UNC) were identified.

### Strand-specific RNA Sequencing

Strand-specific RNA sequencing libraries were prepared using Illumina-compatible NEBNext® Ultra™ II Directional RNA Library Preparation Kit (New England BioLabs, MA, USA) at Genotypic Technology Pvt. Ltd., Bangalore, India. Briefly, 500 ng of total pooled RNA was taken for mRNA isolation, fragmentation, and priming. Two such pools of RNA were sequenced, representing uncomplicated and severe disease complications (denoted with * symbol in Table S1). Fragmented and primed mRNA was further subjected to first-strand synthesis followed by second-strand synthesis. The double-stranded cDNA was purified using JetSeq Beads (Bioline, Cat #BIO-68031). Purified cDNA was end-repaired, adenylated, and ligated to Illumina multiplex barcode adapters as per NEBNext® Ultra™ II Directional RNA Library Prep protocol followed by second strand excision using USER enzyme at 37°C for 15 minutes. Adapter-ligated cDNA was purified and subjected to 11 cycles for Indexing (98°C for 30 sec, cycling (98°C for 10 sec, 65°C for 75 sec) and 65°C for 5 min) to enrich the adapter-ligated fragments. Final PCR products (sequencing libraries) were quantified by a Qubit fluorometer after a quality control check (Thermo Fisher Scientific, MA, USA). The libraries were paired-end sequenced on Illumina HiSeq X Ten sequencer (Illumina, San Diego, USA) for 150 cycles following the manufacturer’s instructions.

### RNA Sequencing Data Analysis

To substantiate the expression profile obtained from microarray analysis, we pooled input RNA from a subset of samples to represent the two populations – severe and uncomplicated malaria and performed directional, paired-end RNAsequencing on these. Hereafter, we refer to this as ‘uncomplicated pool’ and ‘complicated pool’ in the manuscript. Transcriptome analysis was performed by processing the raw data to remove low-quality bases (<Q30) and adapter sequences. The high-quality reads were considered for alignment with the human reference *Homo sapiens* (GRCh38.p13) genome using Hisat2. Unaligned reads against the human genome were mapped against the *Plasmodium falciparum 3D7* reference genome (v64).

Read abundance was calculated for the aligned reads using ***FeatureCounts*** [39] for downstream analysis. For the identification of antisense reads, FeatureCounts was run on the same dataset twice with strand setting reversed, as described by Bao et. al. [40] To identify confident antisense signals, we added a threshold of ≥20 antisense reads in at least one of the sequencing pools. Sequencing (uneven library size/depth) bias among the samples was removed by library normalization using size factor calculation in DESeq2 [41]. Normalized expression values were used to calculate fold change for a given transcript. The differential expression calculation at log2fold change cutoff +/-0.8 is considered. As FeatureCounts would only summarize NATs are read level, we used StringTie (v2.1.2) [42] to assemble transcripts for each gene locus and results were compared with the existing genome annotations (v64) using GFFcompare [43] to summarize NATs at the gene level [44]. To identify antisense splicing junctions, HMMSplicer was used (-p 4 -j 10 -k 3000 -w 4 -e 2 -m 500 -n 700 -d True) as suggested for *Plasmodium* [45–46].

### Functional enrichment by Gene Ontology

Functional enrichment analysis of differentially expressed sense and antisense transcripts in the two disease cohorts was performed individually using DAVID (DAVIDv6.8; http://david.ncifcrf.gov/) against a background list of the consolidated mitochondrial gene list. The GO terms were classified into three categories: Biological process (BP), cellular component (CC), and molecular function (MF), and p<0.05 (EASE score) was considered to indicate a statistically significant difference.

## Results

### Mitochondrial associated genes

A list of 886 genes was prepared which included 42 genes encoded by the mitochondrial genome, 18 encoded by the apicoplast genome, targeted to mitochondria; and 826 encoded in the nucleus and targeted to the mitochondria for their molecular function (Supplementary Sheet1). Out of the total number of genes reported from various prediction tools, 47 genes from TargetP, 463 genes from Cello2GO, 187 genes from MitoFate, 401 genes from PFMpred, and 398 genes from PlasmoDB were shortlisted to be part of the mitochondrial proteome. The criteria used for selection was for a gene to be predicted by at least two of the tools. We further performed literature screening of *P. falciparum* mitochondria associated genes to include them in the study. There appears to be a good overlap between the genes predicted from the literature sources and those from the bioinformatics tools. This comprehensive list of 886 genes (Supplementary Sheet1) therefore comprises 581 genes with published experimental evidence [31–35] and 305 predicted-only genes, and forms the basis for the transcriptomic analyses presented here.

### Identification of sense-antisense transcript pairs specific to disease manifestation

Out of the 886 genes analysed for expression in the CM and HD cohort, 662 genes with sense, and 480 genes with antisense transcripts were found to be expressed in the HD cohort, while 711 sense and 505 antisense transcripts were expressed in the CM cohort. Based on microarray expression data, the differentially expressed genes of CM and HD isolates, compared against those of uncomplicated malaria as control is depicted in Figure 1. There appears to be a much higher number of both sense and antisense transcripts upregulated in the cerebral malaria cohort compared to the hepatic dysfunction disease manifestations. The expression profile of these differentially expressed sense and antisense genes in the individual disease cluster is given in Supplementary Sheet 2.

**Figure 1.**
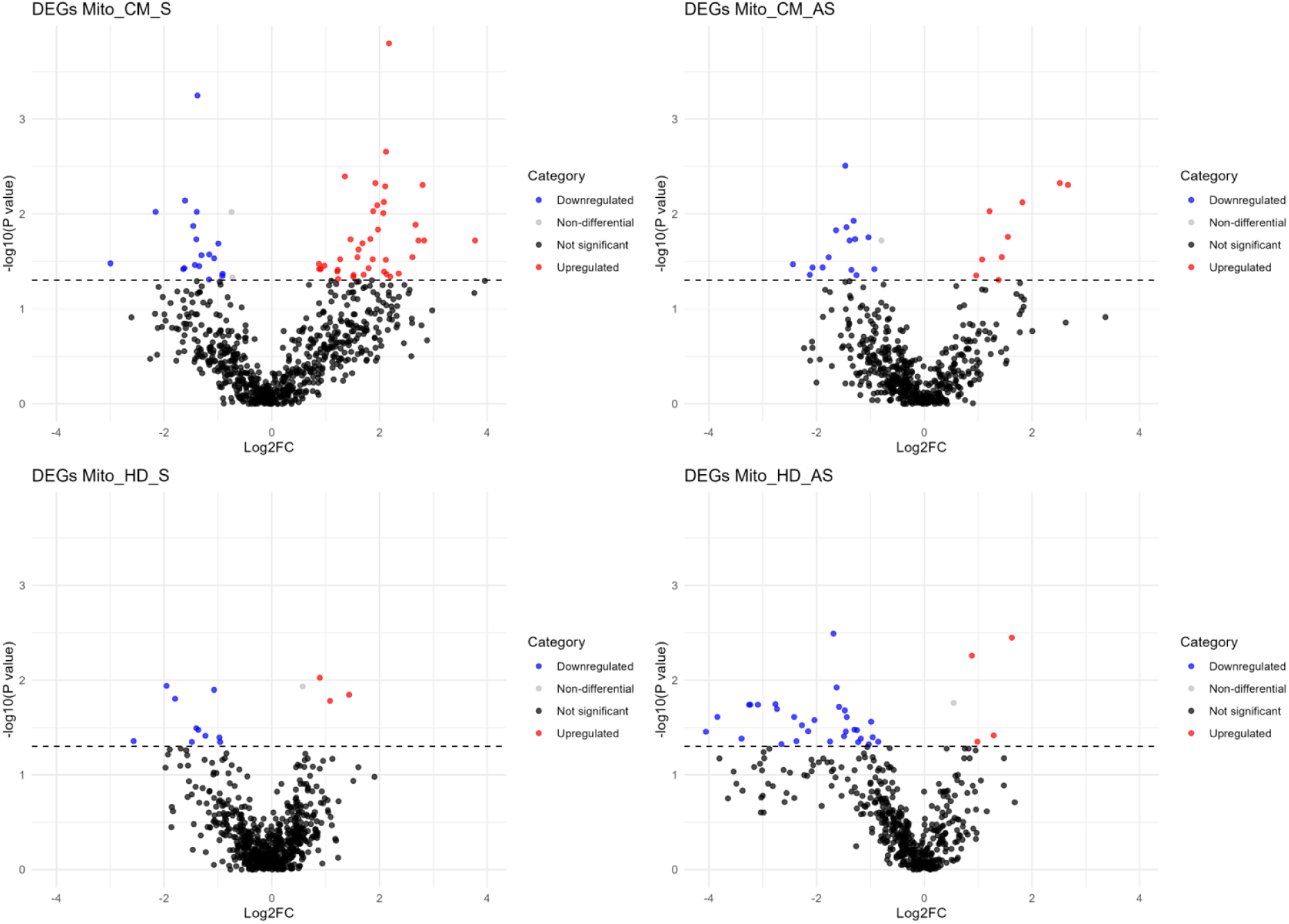
Volcano plot of differentially expressed sense and antisense genes in Cerebral Malaria (CM) and Hepatic Dysfunction (HD) when compared to the uncomplicated malaria cohort, discriminated based on p-value (<0.05) and log2 fold-change value of >=0.8 for upregulated (red dots), <=-0.8 for downregulated (blue dots), and between 0.8 to -0.8 for non-differential gene expression (grey dots).

Out of the 886 shortlisted mitochondria associated genes, 660 sense transcripts, 448 antisense transcripts, and 412 sense-antisense transcript pairs expressed are common between the HD and CM datasets (Figure 2; refer to Supplementary Sheet 3 for transcript IDs). We identified 29 and 59 sense-antisense transcript pairs uniquely expressed in the hepatic dysfunction and cerebral malaria cases, respectively (Supplementary Sheet 3). Much of the sense–antisense transcript pairs in the two disease conditions are positively correlated (70% of S-AS pairs in HD and 50% of S-AS pairs in CM) but there are some genes with a negative correlation between their sense and antisense transcript pairs (0.68% in hepatic dysfunction and 2.54% in cerebral malaria cases), i.e., if sense transcript is upregulated, antisense transcript is downregulated and vice versa. None of the negatively correlated sense-antisense transcript pairs are common between the two disease conditions. A heatmap expression of these transcripts with the gene description is depicted in Figure 3. Except for the gene PF3D7_0923200 (NOS, nitric oxide synthase, putative), all negatively correlated antisense transcripts are downregulated in the HD and CM manifestations.

**Figure 2.**
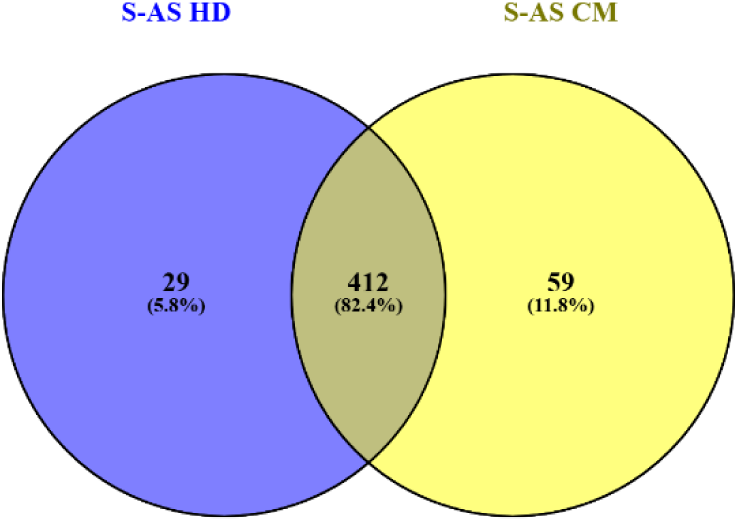
Venn diagram showing the overlap of the number of sense-antisense transcript pairs across two severe manifestations investigated: HD-Hepatic dysfunction, CM-Cerebral Malaria, S-Sense transcripts, AS-Antisense transcripts.

**Figure 3.**
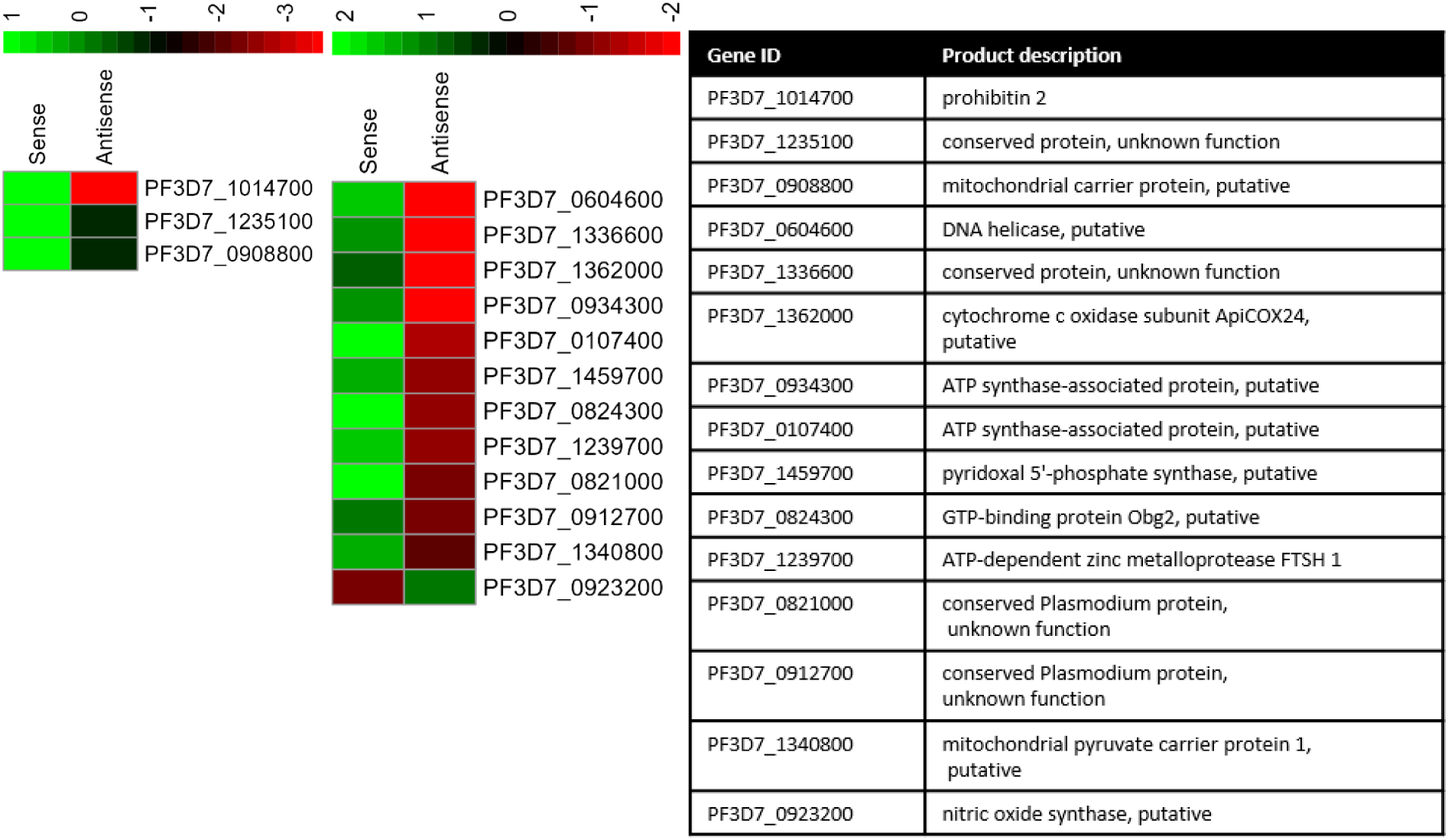
Heatmap diagram showing the gene expression values of sense-antisense transcript pairs that are negatively correlated in the two disease cohorts-HD and CM. individually. The adjoining table corresponds to the description of product these genes as per PlasmoDBv65. HD: Hepatic Dysfunction: CM: Cerebral malaria

### Data validation using RNA sequencing

To validate the gene expression data from microarray experimentation, an RNA sequencing experiment was carried out by pooling 2 severe samples, one each representing the severe HD and CM complication against an uncomplicated pool of sample (marked with * in Table S1). We detected 694 sense and 347 NATs from RNA sequencing based on our filtering thresholds. Comparison of the expression profile of the 536 sense and 139 NATs common between the microarray and RNA sequencing analysis was done by calculating the Pearson correlation coefficient (R) to reflect linearity and concordance between the two experimental approaches, where values closer to 1 depict total positive correlation; -1 depicts total negative correlation; 0 depicts no linear correlation. The correlation plot shows r = 0.83 for the expression of sense transcripts, while antisense expression data shows r= 0.54 (*pval* in both cases <0.05); reflecting a positive correlation between the two experimental gene expression datasets (Figure 4).

**Figure 4.**
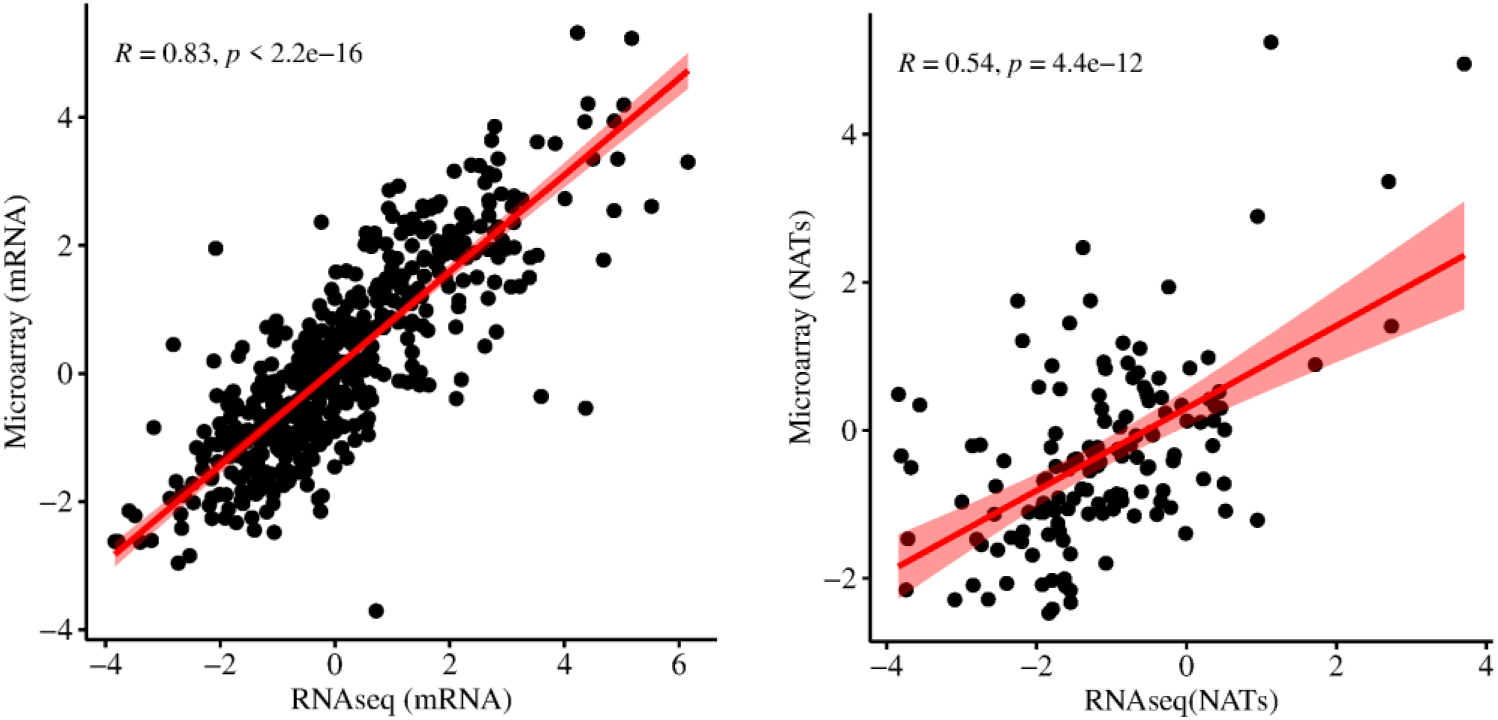
The correlation coefficient (R) for antisense transcripts is 0.54, while that for sense transcripts is 0.83 between microarray and RNA sequencing pool of samples (p-value < 0.05). x-axis denotes the sense and antisense (NATs) transcript expression in RNA seq experiment | y-axis denotes the average fold expression of sense and antisense (NATs) obtained from microarray experiment for the pooled samples used in RNAseq experimentation

Stringtie could assemble antisense transcripts for 15 genes in the uncomplicated and 10 genes in the complicated sample. Their coordinates are listed in Supplementary Sheet 4. For the antisense transcripts assembled in the uncomplicated sample pool, HMMSplicer identified NAT splice junctions for 6 of these genes, out which there was a corresponding junction in the sense transcript for 4 of these transcripts. For the complicated sample pool, there are a total of 4 NATs with at least one splice junction support out of 10 assembled antisense transcripts. Although Stringtie could assemble a fraction of NATs, the substantially higher read density for these loci suggests pervasive, widespread antisense transcription [47–48].

### Gene Ontology-based enrichment of genes with upregulated antisense transcripts

The gene ontology analysis for genes expressing upregulated and downregulated antisense transcripts (pval / EASE score<0.05) in the respective datasets has identified several biological processes unique to individual disease types as depicted in figure 5. In the HD manifestation, GO terms enriched with antisense upregulated genes included intracellular protein transport (GO:0006886) and cellular macromolecule localization (GO:0070727), suggesting a role in intracellular molecular trafficking. Conversely, antisense downregulated genes were significantly enriched in Carbon and pyruvate metabolism (pfa01200 and pfa00620). Key molecular functions included oxidoreductase activity (GO:0016491) and flavin adenine dinucleotide (FAD) binding (GO:0050660). Pathway analysis revealed significant enrichment in TCA cycle (pfa00020), pyruvate metabolism (pfa00620), and biosynthesis of secondary metabolites (pfa01110), indicating reduced role of antisense transcripts in electron transport chain.

**Figure 5.**
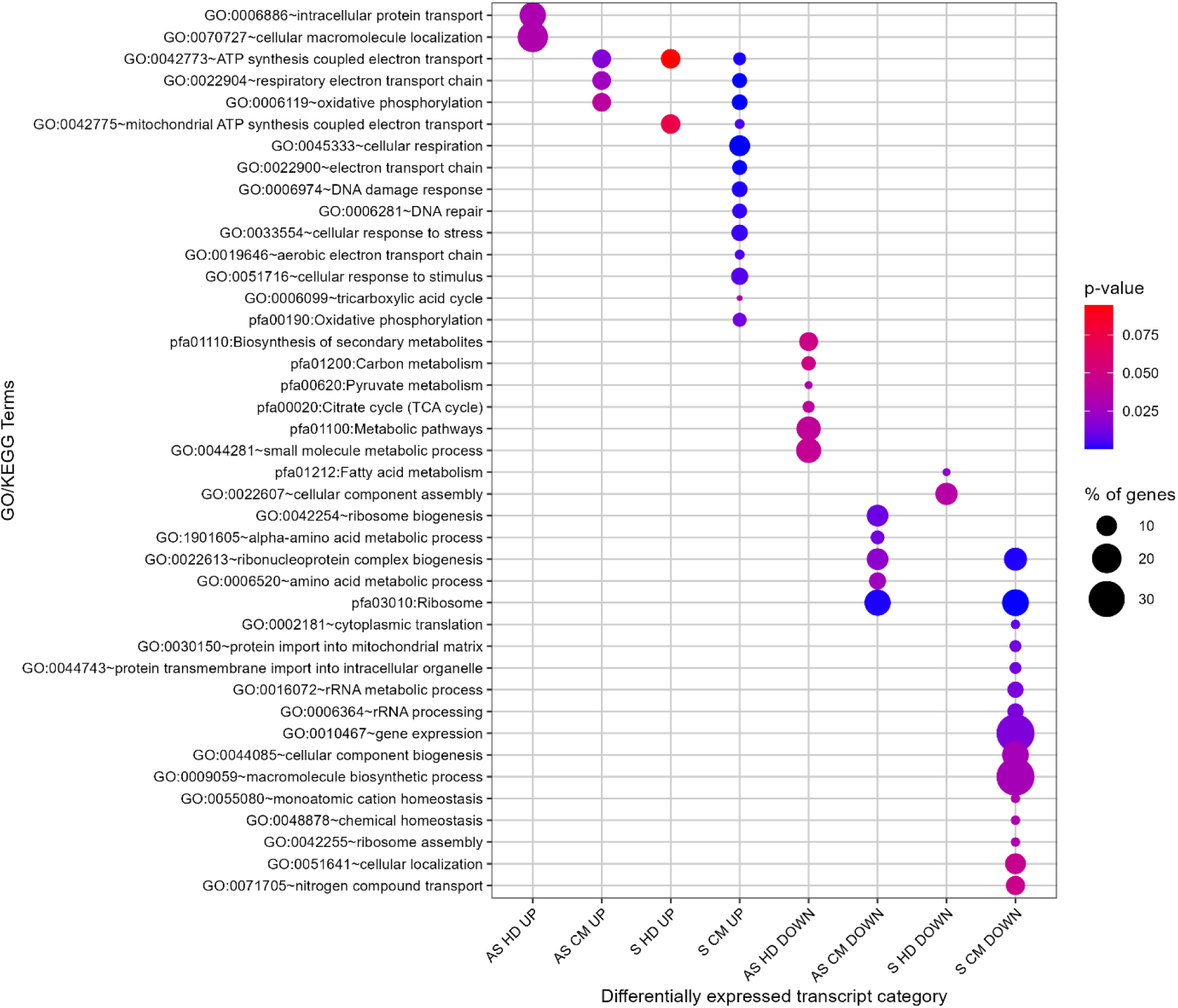
Significant Biological processes and KEGG pathways enriched for differentially expressed sense and antisense transcripts across the two disease types HD: Hepatic Dysfunction: CM: Cerebral malaria (pval <0.05). % of genes denote the fraction of genes in the input list annotated to each GO term. p-value shown are DAVID EASE scores. Enrichment was performed using the study’s consolidated mitochondrial gene list (n=886) as the background.

A similar GO enrichment analysis for the differentially expressed antisense transcripts in the CM cohort show that antisense-upregulated genes were significantly enriched for ATP synthesis coupled electron transport (GO:0042773), and oxidative phosphorylation (GO:0006119). Molecular function enrichment includes DNA binding (GO:0003677) and active transmembrane transporter activity (GO:0022804), suggesting an influence on nuclear processes and membrane transport. Antisense-downregulated genes in this condition were predominantly associated with ribosomal function and RNA processing with the most overrepresented terms being cytosolic ribosome (GO:0022626), ribosome biogenesis (GO:0042254), and RNA binding (GO:0003723). These results suggest a potential suppression of protein synthesis machinery through antisense-mediated downregulation of ribosome-associated genes. The observed differences between the two conditions indicate context-specific expression of antisense transcripts in cellular metabolism and gene expression regulation.

## Discussion

A comparative analysis of 23 species of *Plasmodium* has determined the mitochondrial (mt) genome of the *P. falciparum* to be the smallest [9]. It comprises a 6 kb tandemly arranged, linear, and conserved segment of DNA, encoding only 3 protein-coding genes -*cox1*, (PF3D7_MIT02100), *cox3* (PF3D7_MIT01400) and *cytb* (PF3D7_MIT02300). The remainder encodes several small rRNAs transcribed from both the strands of the mtDNA and ranges from 23-190 nt in length [49]. In total, the mitochondrial genome of *P. falciparum* comprises 42 genes, of which 20 were sufficiently long to be included in the 60-mer probe design in both sense and antisense orientations; the remaining ORFs were too short (<60 nucleotides) for effective probe design and were therefore, excluded from the microarray.

Although the prevalence of NATs encoded from the nuclear genome of the parasite has been well documented [22, 46], the transcription profile of the NATs from the mitochondrial genome of the parasite has been less clear. An overlap of 274 and 285 NATs in HD and CM cluster respectively has been identified after comparing our mitochondrial dataset with the genome wide NATs assembled by Subudhi et. al. [22]. Comparison of our NATs expression data with that of time-based NATs catalogue of laboratory strain of asexual blood stage parasite 3D7 [46] revealed 222 antisense RNAs that appear exclusively in the clinical dataset analysed here (Supplementary Sheet5). Further, comparing the splicing output of HMMSplicer we find splicing evidence of 23 new NATs not reported in the previous study [46], suggesting the parasite actively changing its transcriptome in vivo [50]. Since, PF3D7_1325100 (phosphoribosylpyrophosphate synthetase) shows NAT-associated splice junction in sample pool of both uncomplicated and complicated malaria, this adds an additional layer of transcriptomic complexity that should be taken into account when evaluating the gene as a potential drug target [51].

This study further detected antisense transcripts for all 20 mitochondrial genes included in the array. Upon investigation of the protein coding genes of the mtDNA, we observe an enhanced expression of the sense mRNA of the three respiratory chain-associated genes: *cox1*, *cox3*, and *cytb*, in both the disease manifestations. These three genes form the core components of the parasite’s mitochondrial electron transport chain (mETC), essential for ATP production and parasite survival, and is critical to the parasite transmission from human host to mosquito [49]. The enhanced expression of these genes in the sense orientation (with a corresponding decrease in antisense levels) in severe malaria conditions might reflect the parasite’s increased ATP requirement associated with rapid proliferation, high parasitemia, and the immune-evasive demands of severe host-pathogen interactions during severe malaria infection. Cerebral malaria is often associated with higher parasitemia, increased metabolic needs, and the parasite’s attempt to evade the host immune response [24]; thus, bolstering the efficiency of the ETC could be advantageous for rapid proliferation and survival in this stringent environment. This is further confirmed by the enrichment of biological processes like – *oxidative phosphorylation* and *ATP synthesis coupled electron transport chain* associated with upregulated sense transcripts in CM manifestation. However, further functional studies are needed to confirm this correlation.

The mtETC has been a critical target for malaria treatment [52]. One prominent example is the antimalarial drug atovaquone, which effectively blocks parasite growth by inhibiting the mitochondrial protein cytb during the asexual blood stage of infection [53]. However, resistance against atovaquone has emerged due to point mutations in the cytb gene. Antisense oligonucleotide (ASO)-mediated gene silencing offers a promising alternative therapeutic strategy capable of overcoming such drug resistance by specifically reducing target mRNA expression levels, thus circumventing mutation-based resistance mechanisms [54]. The observed low antisense transcript levels for these critical ETC genes suggest limited transcriptional regulation of these genes by their complementary NATs under severe malaria conditions, potentially facilitating effective ASO-based therapeutic intervention to disrupt parasite proliferation. In addition to mitochondrial encoded ETC genes (*cox1, cox3, and cytb*), nuclear-encoded, apicomplexan-specific ETC subunits (PfCOX6A, PfNDUFA4, PfCOX4, ApiCOX24) also exhibit minimal to no antisense RNA expression, suggesting a conserved pattern of minimal antisense regulation for critical ETC components. This observation underscores the potential for these genes as attractive targets for antisense-based therapeutic approaches, and can be functionally investigated for disrupting essential metabolic processes in malaria parasites.

Despite encoding for three protein-coding genes, plasmodial mitochondria is a validated drug target and has been reported to perform several critical functions for the parasite viz, shuttling electrons for oxidative phosphorylation, iron-sulfur cluster biosynthesis, pyrimidine metabolism, and de novo biosynthesis of heme [8,34]. Owing to its diversified functions, the parasite mitochondria heavily depend on the trafficking of molecules from the nucleus. We have divided these nuclear-encoded transcripts with putative mitochondrial functions into the distinct functional roles and investigated their expression for sense and antisense transcripts across the two disease conditions (Figure 6). The presence of a functional TCA cycle in the blood stages of malaria has remained debatable [9,55]. We have identified several key transcripts of the TCA cycle and mETC, upregulated in cerebral malaria patients (Figure 5). In addition to this, the enrichment of ‘*Oxidative phosphorylation’* (pfa00190) pathway highlights the potential functionality of these genes in driving the parasite’s energy metabolism during blood-stage infection. The genes for the TCA cycle showed expression in parasites in uncomplicated disease conditions. However, they were upregulated in case of cerebral malaria. The identification of upregulated key transcripts of the TCA cycle and mitochondrial ETC in cerebral malaria adds a valuable layer to understanding the parasite’s metabolic adaptations under severe disease conditions. Studying the protein abundance of these genes would provide conclusive evidence of the extent to which the parasite brings its oxidative phosphorylation into effect in the vertebrate host.

**Figure 6.**
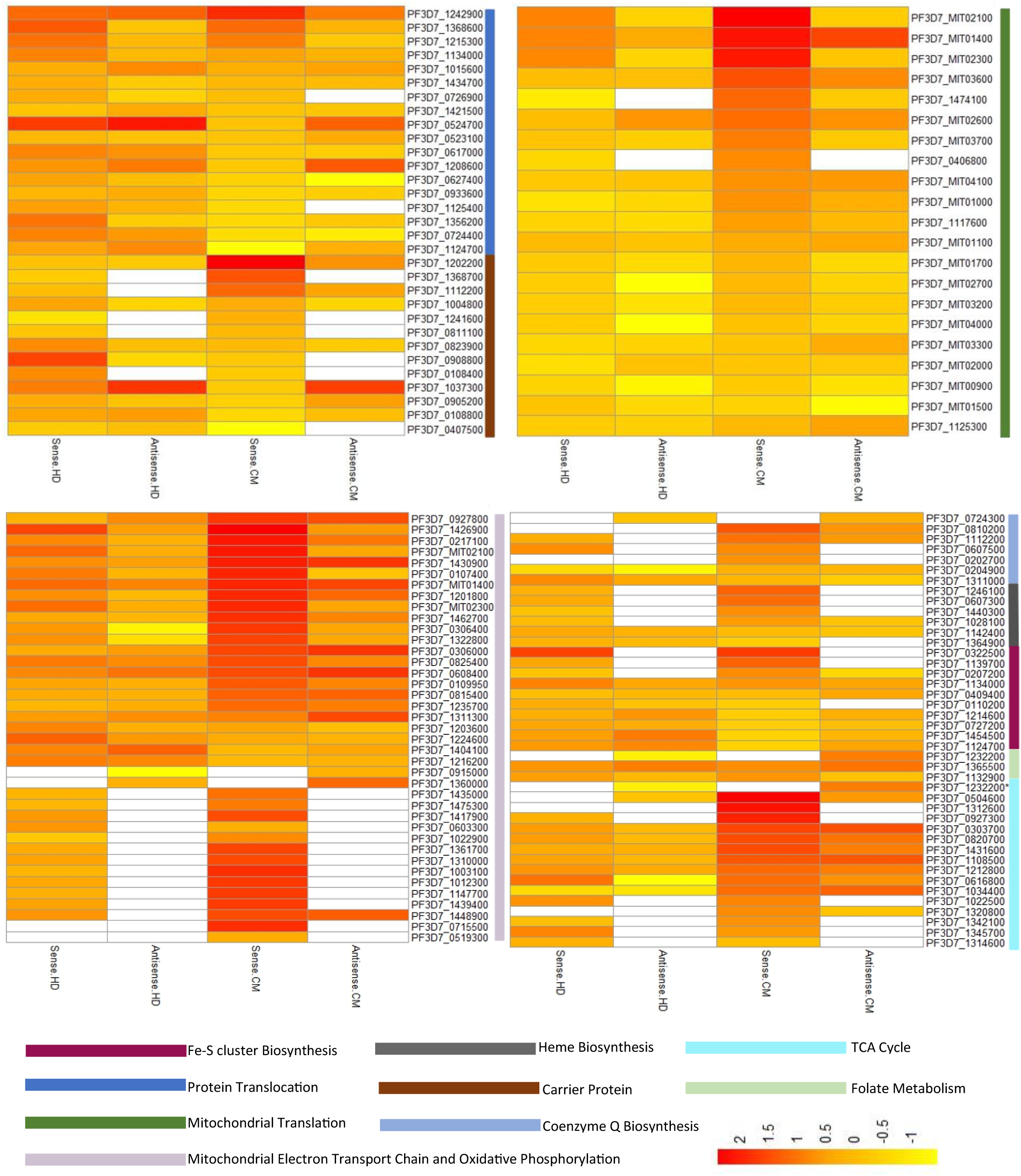
Heatmap expression of sense and antisense transcripts. The transcripts are categorized by their functions, which are indicated by the colored tiles on the left side of the heatmap. White represents no transcript detected corresponding to that gene for the disease group. HD: Hepatic Dysfunction: CM: Cerebral malaria.

*Plasmodium* mitochondria encode 27 rRNA ‘minigenes’ which form the structural component of the parasite’s mitochondrial ribosome (mitoribosome), enabling mitochondrial protein synthesis to occur [7, 56]. The pivotal role of mitoribosomes has been characterized in translating genes encoded in the mtDNA, and a conditional knockdown of these transcripts has shown reduced activity of the cytochrome bc1 complex [57]. Our expression dataset has detected the prevalence of 12 ncRNA genes encoding for ribosomal rRNA fragment in both sense and antisense orientation in all the clinical isolates hybridized under investigation. The inability to detect the expression of the remaining ncRNAs could be attributed to experimental limitations of microarray and short-read RNAseq and emerging technologies employing longer-read RNAseq could help capture the complete transcriptome for these ncRNAs [58]. However, the expression of both sense and antisense transcripts for the genes involved in mitoribosome assembly across both complicated and uncomplicated disease states warrants the fact that the *Plasmodium* mitochondria have a functionally active protein translational machinery in asexual ring stage of development. Another molecule, ICT1 (PF3D7_1117600), that is perpetually expressed across all disease states, encodes for peptidyl-tRNA hydrolase enzyme to catalyze hydrolytic cleavage of prematurely terminated peptidyl-tRNA moieties in stalled mitoribosomes from the mitochondrial ORFs of humans [59–60]. Its critical role in gene expression in organellar translation identifies this as an important antimalarial target [61], warranting further investigation. Transcriptional interference of these ribosomal RNAs by complementation with their respective antisense RNAs could be a plausible mechanism to control gene regulation in the parasite by stalling mitoribosome assembly. However, further experimental validation is necessary to confirm this regulatory model.

Mitochondrial gene regulation is increasingly recognized as an important contributor to the cell physiology [62–63]. NATs have been recognized for their roles in controlling gene expression at multiple levels, including transcriptional interference and chromatin remodeling [64–66]. In the context of malaria disease biology, these patterns of sense/antisense expression in the parasite’s mitochondrion may have functional significance. We have observed discordant (high sense and low antisense) gene expression of fifteen S-AS transcripts in the two clinical manifestations investigated (Figure 2). Four of these (PF3D7_1235100, PF3D7_1336600, PF3D7_0821000, PF3D7_0912700) encode for conserved *Plasmodium* proteins and have not been characterized for being associated with the *Plasmodium* mitochondria, making them important targets for functional assays.

While most of these negatively regulated gene pairs show high sense and low antisense transcription, we observed that only one gene, NOS (PF3D7_0923200, nitric oxide synthase), exhibited inverse expression pattern with elevated levels of antisense transcripts alongside reduced levels of its corresponding sense transcript. This pattern suggests the potential involvement of antisense RNA in the negative regulation of NOS expression, thereby modulating nitric oxide (NO) production. Low NO levels have been associated with more severe disease [67–68]. In an experimental cerebral malaria model, inhaled nitric oxide reduced inflammation and improved survival of the infected mice [69]. Given that host-derived NO exerts antiparasitic effects [70], the parasite’s own control over NO synthase expression could be critical. If the parasite were to produce large amounts of NO through its own NOS activity, this might be detrimental to its survival. In this context, antisense regulation could serve as a safeguard, maintaining NO production at levels that are not self-damaging to the parasite. This finding supports previous reports that highlight the importance of finely tuning NO production during malaria infections [71].

We have further presented a comprehensive list of upregulated antisense transcripts in individual disease manifestations (Supplementary Sheet 2). Four of them are encoded by the nucleus (PF3D7_0410800, AKIT3; PF3D7_0524700, TOM22; PF3D7_0719100, ATP synthase F0 subunit a-like protein and PF3D7_1037300, AAC1) and consistently exhibit elevated antisense transcription in both the disease manifestations. These genes are critical components of the mitochondrial protein import machinery in *P. falciparum*. Specifically, AAC1 (PF3D7_1037300 -ADP/ATP carrier protein 1) which functions as an ATP/ADP antiporter, plays an essential role in the energy metabolism of the parasite [72]. Transcriptomic profiling of antisense genes from culture strain of *P. falciparum* detected low abundance of this gene across all four time points [46] in contrast to the transcriptome reported in the current study. This discrepancy emphasizes the need for both in vitro and ex vivo datasets for a comprehensive and accurate understanding of parasite gene expression. TOM22 (PF3D7_0524700) is another key gene, encoding the mitochondrial import receptor subunit TOM22, which is translocated to the outer membrane of mitochondria for the assembly of the TOM complex (translocase of the outer membrane of mitochondria) for importing mitochondrial proteins [72–73]. Collectively, these results support the idea that mitochondrial transport genes are promising targets for further investigation in severe *P. falciparum* malaria, for several reasons: i) their functional assembly mediates the uptake of solutes essential for parasite proliferation and survival; hence, perturbing their function can compromise critical mitochondrial processes; (ii) *P. falciparum* exhibits resistance against various conventional transporters [74], signalling the need for new therapeutic strategies less prone to resistance; and (iii) the consistent abundance of antisense transcripts for these genes in both the disease clusters may suggest their regulation under control by sense/antisense interaction. Further, direct human homolog of these genes shows negligible similarity by standard BLAST searches, enhancing the clinical potential of these targets. A functionally analogous receptor (TOMM22) does exist in humans, but does not share high sequence identity. The upregulated NATs associated with these genes may form part of a network that ensures timely production and assembly of the corresponding protein-coding transcripts [75–76], serving as important control points in parasite biology. Elevated NATs can significantly perturb the translational potential of their targets [71, 75–77]. Future work involving targeted knockdowns, CRISPR interference, or other functional approaches against these genes is warranted to validate their precise regulatory roles and clinical relevance as potential biomarkers in severe malaria.

Studies on non-coding RNAs of *P. falciparum* have only recently been taken up by multiple groups [46, 66, 78]. A constant challenge in understanding parasite disease biology is that most of these studies work in culture systems. Building upon our previous profiling of natural antisense transcripts (NATs) from clinical *P. falciparum* datasets, the present work provides the first comprehensive transcriptomic dataset of parasite mitochondria from 22 clinical isolates, capturing their differential expression across two of the most severe malaria complications [22]. Since the differences in parasite developmental stage has been a confounding factor in studies characterizing clinical transcriptome, we attempted to estimate the stage of the mitochondria-associated transcriptome in this study by comparing our gene expression dataset to that of previously published high-resolution IDC transcriptome [79] individually for the CM and HD cohort. We find that the expression profile of the majority of genes map to trophozoite (61%) and late ring/early troph stage (16%) for both the disease cohorts. Although this does not replace stage correction, it indicates broadly similar stage distributions across cohorts. Moreover, the precise time point/staging of maximum gene expression in a clinical sample might shift under host pressure in different disease states [80], making sample heterogeneity a limiting factor in clinical studies.

Given the minimal role of mitochondria in blood-stage malaria [9], the extensive presence of these NATs from mitochondrial ORFs in severe malaria patient isolates may indicate their regulatory functions in parasite biology. Since the mitochondrial genome is arranged in tandem repeats, detecting both sense and antisense transcripts hints at active post-transcriptional regulation in the mitochondrion. Despite inherent challenges such as uneven cohort sizes, limited RNA-seq representation, and variations in parasite stage and parasitemia, our findings substantially extend the sparse field of *Plasmodium* mitochondrial transcriptomics, especially from clinical samples. Looking ahead, the adoption of long-read sequencing technologies will be instrumental in resolving the full diversity of mitochondrial ncRNAs, while functional approaches (e.g., antisense oligonucleotides and CRISPRi-based perturbations) will be essential to validate sense–antisense regulatory mechanisms. While our study’s scope is limited by its single-center cohort and a modest sample size, the rarity of severe malaria cases and the sparsely characterized mitochondrial transcriptome from field isolates of *Plasmodium* make this work a valuable resource for guiding future studies targeting mitochondrial gene regulation in *Plasmodium* infection. This work expands our understanding of *Plasmodium* mitochondrial biology, highlights the interplay of these sense and antisense transcripts in the context of mitochondrial microenvironment of the parasite under host pressure, emphasizing key genes whose investigation could provide us with potential targets for novel therapeutic interventions against *P. falciparum* malaria.

## Acknowledgments

We thank all the patients and technical workers for participating in and supporting this project. This work is funded by the Indian Council of Medical Research (ICMR) under the Extramural Ad-hoc Scheme (PID: 2019-1121, a multicentric project with AD as PI and DG and SKK as Co-PI). S.G. was supported by ICMR as project assistants to carry out this work and was subsequently supported by Institute fellowships for the PhD program applicable by BITS Pilani (Pilani campus). We thank Birla Institute of Technology (BITS), Pilani (Pilani campus), International Centre for Genetic Engineering and Biotechnology (ICGEB), New Delhi, and SP Medical College, Bikaner, for providing the necessary facilities required for this work. We also thank Genotypic Technology Pvt. Ltd., Bangalore, India, for the microarray hybridization and RNA sequencing experimentation and consultation regarding data analysis.

## Availability of data and materials

All microarray results have been deposited in the Gene Expression Omnibus database (GSE272963). Raw sequencing data has been uploaded on IBDC database under controlled access with accession no. INRP000336 (Bioproject Accession: PRJEB90534) along with metadata. The INSDC sample accession IDs are ERS25049603 and ERS25049604 for complicated and uncomplicated sequencing experiments, respectively. All patient metadata are included in Supplementary Sheet1 (Table S1). Reprint requests should be addressed to AD.

## Ethics statement

Informed consent was obtained from all individual participants in the study according to the SP Medical College Hospital’s guidelines (No. F (Acad)SPMC/2003/2395). Permission to use these samples for further studies was given through IERB approval No. F29(Acad)SPMC/2020/3151 dated 05.09.2020 and fresh samples were collected according to EC/SPMC/ECA/06/28-02-2019. We thank all the patients for their voluntary informed consent and participation.

## Author Contributions

AD conceived the study, acquired funding from ICMR, India, and networked to facilitate the outcome. SG was involved in performing the experimentation, analysis, and data interpretation, designing the figures, and drafting the manuscript under the constant guidance of AD. BS assisted in sequencing analysis. DG has been closely involved in consultations on data analysis and related discussions since the study’s inception. DKK and SKK supervised sample collection and clinical characterization. All authors read and approved the manuscript.

